# Adaptation to transients disrupts spatial coherence in binocular rivalry

**DOI:** 10.1101/867374

**Authors:** Marnix Naber, Sjoerd Stuit, Yentl de Kloe, Stefan Van der Stigchel, Chris L.E. Paffen

**Author notes:** **Corresponding Author** Dr. Marnix Naber, Room H0.25, Heidelberglaan 1, 3584CS Utrecht, The Netherlands.

## Abstract

When the two eyes are presented with incompatible images, the visual system fails to create a single, fused, coherent percept. Instead, it creates an ongoing alternation between each eye’s image; a phenomenon dubbed binocular rivalry (BR). Such alternations in awareness are separated by brief, intermediate states during which a spatially mixed (incoherent) pattern of both images is perceived. A recent study proposed that the precedence of mixed percepts positively correlates with the degree of adaptation to conflict between the eyes. However, it neglected the role of visual transients, which covaried with the degree of conflict in the stimulus design. We here study whether the presence of visual transients drive adaptation to interocular conflict and explain incidence rates of spatially incoherent BR. Across three experiments we created several adaptation conditions in which we systematically varied the frequency of transients and the degree of conflict between the eyes. Transients consisted of grating orientation reversals, blanks, and plaids. The results showed that the pattern of variations in the fractions mixed percepts across conditions was best explained by variations in the frequency of visual transients, rather than the degree of conflict between the eyes. We propose that the prolonged presentation of transients to both eyes evokes a chain of events consisting of (1) the exogenous allocation of attention to both images, (2) the increase in perceptual dominance of both rivalling images, (3) the speed up of adaptation of interocular suppression, and eventually (4) the facilitation of mixed perception during BR after adaptation.

**Author summary:** When one eye is presented with an image that is distinct from the image presented to the other eye, the eyes start to rival and suppress each other’s image. Binocular rivalry leads to perceptual alternations between the images of each eye, during which only one of the images is perceived at a time. However, when the eyes exert weak and shallow mutual suppression, observers tend to perceive both images intermixed more often. Here we designed an experiment and a model to investigate how stereoscopic stimuli can be designed to alter the degree of interocular suppression. We find that prolonged and repeated observations of strong visual transients, such as sudden changes in contrast, can facilitate the adaptation to suppression between the eyes, resulting in that observers report more mixed percepts. This novel finding is relevant to virtual- and augmented reality for which it is crucial to design stereoscopic environments in which binocular rivalry is limited.

## 1 Introduction

### 1.1 Studying the dynamics of visual awareness with binocular rivalry

Binocular rivalry (BR) is a primary method in the scientific fields of cognitive psychology and neurosciences to study visual awareness. It consists of the presentation of separate images to each eye. When the two images are distinct, the visual system is unable to fuse them into a coherent percept. Instead, the distinct mental representations of both eyes compete for priority to visual awareness. This results in the perception of unending perceptual alternations between the two images over time, a purely internally (mentally) driven process because the physical environment is kept stable.

BR has been heavily exploited by psychologists, neuroscientists, and philosophers for a variety of reasons. One reason is that the dynamic properties of BR provide information on what type of images dominate more strongly or break into visual awareness faster (e.g., 1, 2). Such research is necessary in order to understand why people sometimes fail to notice objects (e.g., in traffic), how image-parts are grouped into ensemble objects (i.e., Gestalt principles), and why certain objects in the visual environment receive sensory priority (e.g., advertisements). BR is also the primary method used to study the interaction between the sensory processing of stimulus properties and other cognitive high-level functions such as attention, numerosity and emotions (3–7). Furthermore, studies have revealed that a variety of brain regions and processes underlie changes in the content of visual awareness during BR (8–11). Using BR to find the neural loci of consciousness and to identify the distinct processing stages of the stream of consciousness remains an ongoing line of research. Lastly, BR serves as a tool to examine to what degree information, that falls outside the scope of awareness, is processed and affects behavior (e.g., 12). Following the iceberg-mind analogy in the sense that most of what an iceberg’s constitutes is submerged under water, most stimuli in a visual environment are not consciously perceived but may still have a determinative effect on decision-making (13). In sum, BR has been shown to be a valuable method to examine perceptual selection, the neurobiological underpinnings of awareness, and unconscious processing (14). However, there is more to be learned from BR. While often overshadowed by discussions surrounding consciousness, BR also reflects how the eyes interact and strive for a stable, coherent percept. It is therefore necessary to understand under which circumstances dichoptic images fuse and when they engage in binocular rivalry (e.g., 15, 16–18). Especially now, with the rise of virtual and augmented reality goggles, it is of importance to understand how images can be best designed to prevent BR, enhance the fusion of representations of both eyes, and create realistic depth perception. The experience of a coherent percept is important for effort-free viewing and the feeling of immersion when wearing stereoscopic goggles (19). BR may thus also be utilized to determine the level of “cooperation” between the eyes.

### 1.2 Exclusive versus nonexclusive, mixed episodes in binocular rivalry

How can BR serve as a tool to determine to what degree information of both eyes integrates rather than competes? To answer this question, it helps to focus on the spatio-temporal dynamics rather than merely the temporal dynamics of BR. Temporal dynamics include the rate at which switches in awareness occur and the ratio of left versus right eye dominance durations in perception. These measures indicate when and how often a change in awareness occurs and how strong, conspicuous, and relevant each image is to the visual system. Spatio-temporal dynamics embrace the local nature of binocular conflict and include episodes in which BR is in an intermediate, unstable state, in which perception exists of a mixture of the images of both eyes across image locations (i.e., piecemeal or non-exclusive rivalry). This latter measure indicates to what degree information of both eyes is integrated. However, this aspect of BR has received relatively little scientific attention, mainly because the spatio-temporal dynamics of binocular rivalry are typically operationalized as stemming from a discrete on-off process (i.e., the image of the left or right eye is visible) by means of measurements of binary responses (i.e., press either one of two buttons to report dominance of the two images). Only a handful of papers have looked at the non-binary properties of rivalry. For example, Naber et al. (20) instructed observers to report mixed percept episodes of moving gratings with a joystick and observed that the reported spatio-temporal dynamics matched the same dynamics measured objectively with the optokinetic nystagmus. Other studies examined so-called traveling dominance waves, described best as the gradual emergence of a suppressed image as it flows over the other, dominant image within a relatively short time span (21–23). These waves tend to have a local starting point in the visual field and move with a certain velocity (24–29). A few more studies inspected what type of images proliferate mixed percepts during rivalry (30–32). For example, the more similar the images are across the eyes, the weaker the interocular suppression and the higher the chance of observing mixed rivalry (31). Similarly, gratings which are relatively similar with locally overlapping features, exhibit more mixed percepts as compared to complex, coherent objects such as houses and faces, which are more dissimilar (32). This means that when images mutually exert weak, shallow interocular suppression (i.e., a weak competition between the eyes) due to a local overlap of features between both eyes, exclusive (monocular) percepts are rarer and mixed episodes last longer. A recent adaptation study additionally showed that the durations of mixed episodes can be lengthened by first adapting observers to episodes of strong interocular conflict in orientation (33). The explanation for this finding is that the visual system includes neurons that detect conflict between the eyes and drive interocular suppression (34). When these conflict detectors become less responsive due to adaptation, interocular suppression presumably becomes weak (i.e., shallow), resulting in more or longer episodes of mixed rivalry. However, adaptation to interocular conflict may not be the only plausible explanation for the reported effects on mixed percepts during rivalry. The current study investigates whether the weaker suppression (reflected by a larger incidence of mixed percepts) following adaptation in the study of Said and Heeger was due to adaptation to conflict, or whether other factors contribute to weaker suppression following adaptation.

### 1.3 Binocular conflict detectors versus visual transients

Although Said & Heeger (33) elegantly applied the method of adaptation to support their model including conflict detectors, the authors may have overlooked the possibility that additional or alternative mechanisms may drive the occurrence of mixed episodes. Here we propose that the presence of strong transients affects binocular rivalry and, in the context of the findings of Said and Heeger, could be the principal underlying factor for the facilitation of mixed percepts in rivalry after adaptation. To clarify, let us first describe how visual transients affect binocular rivalry: It is known that an intermittent stimulus presentation (i.e., interleaving content-rich image presentations with content-absent blanks) strongly reduces the alternation rate of binocular rivalry. Depending on the duration of the blank episodes, an image of one eye can remain dominant for minutes rather than seconds (35, 36). Such changes in the temporal domain of rivalry dynamics suggest that intermittent presentation enhances interocular suppression. We here propose that intermittent presentation (i.e., a strong visual transient) also affects the spatio-temporal rivalry dynamics. As for the study of Said & Heeger (33), their conflict condition (producing strong adaptation) included an intermittent presentation paradigm while their weak adaptation condition did not (see Figure 1, a modification of Figure 6 in Said & Heeger). In other words, the implementation of blanks, and thus of transient onsets and offsets of the images, may have facilitated adaptation to interocular suppression rather than conflict.

**Figure 1.**
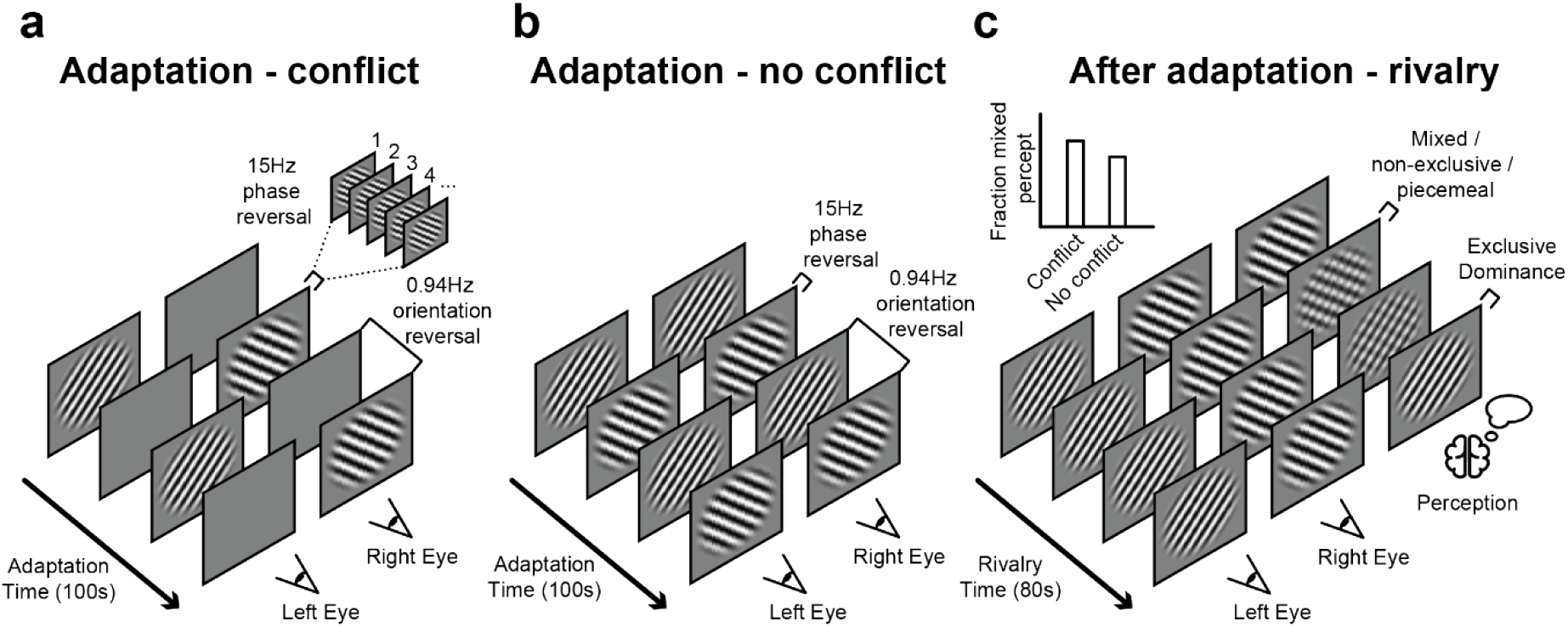
Procedural design by Said & Heeger. In the design of Said & Heeger’s (33), a single trial consisted of an adaptation period (a-b) that lasted for 100s and a rivalry test period (c) that lasted 80s. Observers experienced regular binocular rivalry and indicated the onsets of exclusive and non-exclusive percepts with keyboard buttons during the subsequent test period (c). During the preceding adaptation period, observers passively viewed alternations in oriented gratings (a-b). The gratings’ phase reversed at a rate of 15Hz to prevent local brightness adaptation. More importantly, the perceived orientation alternated (counter-)clockwise at a rate of 0.94 times per second. According to Said & Heeger (33), prolonged presentation of different orientations to the eyes (a) adapt opponency neurons that detect interocular conflict and drive interocular suppression (i.e., the degree the left eye’s image is inhibited by the right eye’s image and vice versa). However, when identical orientations are presented to both eyes at any point in time (b), they argue that no conflict between the eyes is present and neural conflict detectors do not adapt. Adaptation and therewith weaker interocular suppression subsequently leads to unstable perception, that is, a higher precedence of mixed (nonexclusive, piecemeal) percepts. A mixed percept consists of the presence of parts of two images from both eyes rather than a single exclusive image of one eye (c).

In three separate experiments we demonstrate that the presence of visual transients during adaptation explains the degree of mixed percepts better than the presence of orientation conflict between the eyes. By manipulating the rate of changes in monocular contrast and changes in orientations, we are able to show that these transients affect interocular suppression, resulting in decreased spatio-temporal stability of binocular rivalry.

## 2. Experiment 1

### 2.1 Introduction

As described in the introduction, incoherent rivalry after adaptation (i.e., more mixed percepts) may be caused by the presence of transients, rather than the presence of interocular conflict. Such transients can be of any type, including changes in contrast and orientation. Here we extended the original conflict and no conflict conditions of Said & Heeger with novel conditions that either included or excluded the different transient types described above (see Figure 2a; for details, see *Stimulus and conditions*), and investigated their individual contributions to the degree of mixed percepts following adaptation.

**Figure 2.**
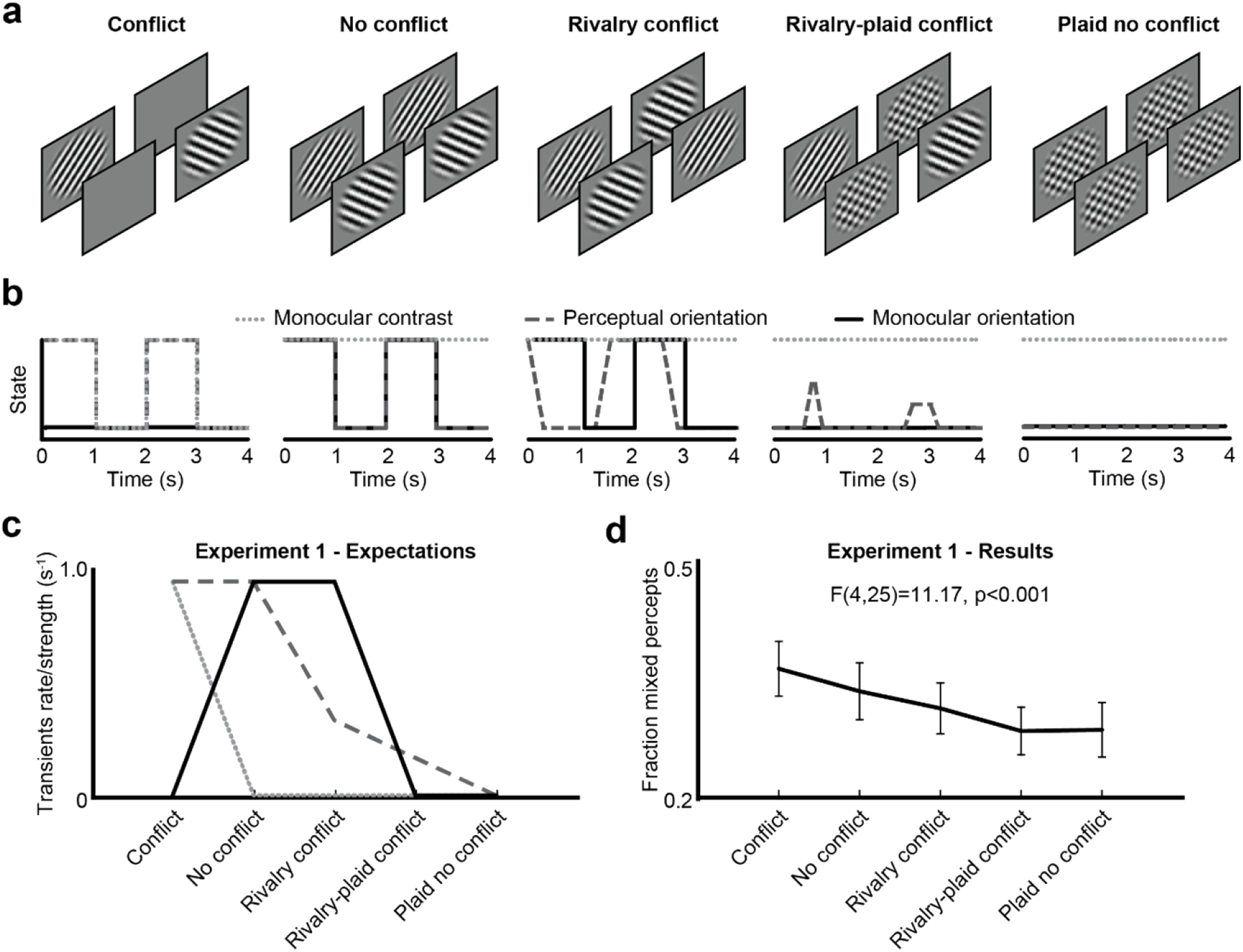
Adaptation conditions, transient profiles, predictions, and results of Experiment 1. Experiment 1 tested five adaptation conditions with different stimuli (a). Each condition induced changes in stimulus states as a function of time (b), including monocular (solid black) and perceptual orientation (dashed dark gray) transients, and contrast transients (dotted light gray). We included the original conditions of Said & Heeger in which orientation and contrast (first panel from left) or only orientation (second panel) changes at a rate of 0.94Hz (solid black lines at (b)). We extended the original design by including a rivalry condition (third panel) with less frequently perceived orientation reversals (dashed dark gray lines at (b)) and a continuous orientation conflict between the eyes that excludes a monocular contrast conflict (dotted light gray lines at (b)). Another condition similar to the first panel was added but with a plaid rather than a blank screen in the other eye (fourth panel). A fifth condition with plaids presented to both eyes and thus no transients served as a baseline. Note that the perceptual orientation transients (dashed dark gray) are visible to the observer while the monocular transients (dotted light gray and solid black) are not (b). The pattern of expected fraction mixed percepts across conditions per feature (c) is based on the number and strength of transients within a normalized time interval (for legend, see panel b). The actual pattern of fraction mixed percepts as indicated by observers (d) did not perfectly match the patterns predicted by each individual adaptation type but matched a combination of monocular contrast and perceptual orientation factors (c).

### 2.2 Methods

#### 2.2.1 Participants

Twenty-six human individuals, all right-handed, young students (age: *M =* 23.4, *SD =* 4.5; 21 females) and with normal or corrected-to-normal vision, participated in Experiment 1. Participants were naïve to the purpose of the experiment, gave informed written consent before participation, and received either study credit or money (€6 per hour; Experiment lasted approximately 3 hours) after participation. The experiments conformed to the ethical principles of the Declaration of Helsinki and were approved by the local ethical committee of Utrecht University.

#### 2.2.2 Apparatus

Stimuli were generated on two 24-inch ASUS VG248QE monitors (AsusTek, Taipei, Taiwan) with a dell computer (Dell, Round Rock, TX, USA) operating Windows 7 (Microsoft, Redmond, WA, USA) and MatLab version r2010a (Mathworks, Natick, MA, USA). The presentation monitors displayed 1920 by 1080 pixels at a 60-Hz refresh rate. Each screen size was 53cm in width and 30cm in height (51 by 29 visual degrees), and the participant’s viewing distance to the screen was fixed with a chin and forehead rest at 57cm. Each eye of an observer was presented with stimuli through a Wheatstone-inspired (37) mirror stereoscope (for details, see (38). Observers used the arrow buttons on a Logitech keyboard (Logitech International S.A., Lausanne, Swiss) to report their percept (left for exclusive dominance of counter-clockwise-oriented gratings, down for non-exclusive dominance, and right for exclusive dominance of clockwise-orientated gratings).

#### 2.2.3 Stimuli and conditions

We used stimuli and conditions similar to those of Said & Heeger (33) by including an adaptation phase (Figure 1a-b) to affect perceptual stability in a subsequent rivalry phase (Figure 1c). Stimuli had a 0.6° radius in visual angle, a spatial frequency of 6.6 cycles/°, and edges softened by a cosine ramp of 0.1° in width. To prevent ocular vergence responses and thus to promote binocular fusion (i.e., to achieve perception of two spatially overlapping images), the stimuli were surrounded by a fusion stimulus. The fusion stimulus consisted of a 0.3° wide annulus (not shown in the figures) with a random noise pattern that was identical for each eye’s image and located at 2.25° eccentricity.

Besides incorporating Said and Heeger’s two original adaptation conditions in our design (first two panels from the left in Figure 2a), we added an adaptation condition called “rivalry conflict” (panel three in Figure 2a). In contrast to the conflict condition (panel one in Figure 2a), this condition did not include monocular contrast transients (i) but did include monocular orientation transients (ii) and could potentially adapt opponency neurons due to the conflicting information between the eyes.

Note that both the conflict and no conflict condition produce clearly visible transients in orientation at a fixed rate of approximately one reversal per second. However, the rivalry conflict condition also induces alternations between the eyes at a rate dependent on the perception of the observer. This condition thus adds another research opportunity, namely to investigate to what degree the rate of perceived, binocular transients affect the stability of binocular rivalry. Therefore, to manipulate and weaken conflict between the eyes even further in an incremental manner, and therewith the rate of perceived orientation alternations, we added two more conditions with plaids (see panel four and five in Figure 2a). These conditions serve as a baseline in which hardly any transients in terms of contrast and orientation are produced.

While the rate of *physical* stimulus changes was kept constant at 0.94Hz in the original two conflict and no conflict conditions, the three novel conditions were expected to differ in the number of evoked *perceptual* changes in orientation. Specifically, the rivalry conflict condition (third panel in Figure 2a) should evoke perceptual rivalry as probed in the test phase. The rivalry-plaid conflict condition (fourth panel) should cause even fewer perceptual reversals because the images of the plaid and the oriented grating are typically merged in a single percept during binocular rivalry (16). Note that the rivalry-plaid condition was similar to the original conflict condition of Said and Heeger but included the presentation of a plaid rather than a blank screen to the other eye as the oriented grating. The plaid no conflict condition (fifth panel) should cause no rivalry (16, 33). As the observers did not report alternation rates during the adaptation phase, authors MN and YdK independently confirmed that the orientation reversal rate and perceptual appearances were indeed manipulated as intended. In addition to the perceptual orientation transients, the five conditions also differed with regard to the presence of monocular contrast transients. Only the original conflict condition included intermittent blank presentations. The other four conditions thus contained no contrast transients (i.e., second-order, nonluminance contrast).

As shown in Figure 2b, the frequency and strength of each type of transient should differ considerably across the five conditions. For each transient type we plotted a hypothetical pattern of results (Figure 2c) assuming that each specific transient type independently affected the fraction mixed percepts during the test phase. Later in this paper we modelled weighted combinations of multiple transient types to investigate which of these best explain the fraction across all conditions and experiments (see last result section).

#### 2.2.4 Procedure

The task for an observer was to attentively view the stimuli during the adaptation phase. Next they indicated their percept during the binocular rivalry phase as either exclusive (i.e., the majority of the surface of a single image was dominant) or mixed. The observers knew when to start reporting perceptions because the start of the rivalry test phase was marked by a sudden offset of phase reversals (i.e., the stimuli were contrast reversed at a rate of 15Hz during the adaptation phase to prevent local brightness adaptation; see Figure 1a-b). The observers kept their gaze on the fixation point at the center of the stimuli and screen.

Each condition was tested with six trials. The conditions were counterbalanced and the trials were divided into two experimental sessions held at different days, because the experiment took more than 3 hours in total. Both sessions of the experiment started by having the observers align the stimuli on the screens to achieve best fusion, that is, the observers made sure the rivaling stimuli overlapped when viewed through the mirror stereoscope. Next, observers performed one rivalry test trial during which the contrast of the gratings was adjusted with the goal to counterbalance eye dominance by annulling between eye differences in dominance durations. Next, participants performed 30 trials and initiated the start of each trial with a button press.

#### 2.2.5 Analysis

We refer to the independent variable as *adaptation type*. The dependent variable was the *fraction mixed percepts*, that is, the fraction of the total duration of perceptual episodes consisting of mixed dominance, as indicated by the observer. To investigate whether the factor adaptation type significantly affected the fraction mixed percept, we conducted a repeated measures ANOVA as a statistical test of significance (see figures for statistical outcomes). As post-hoc tests, we compared the fraction mixed percepts between each possible pair of conditions with two-tailed dependent t-tests (see tables in supplementary materials for statistical outcomes). We also examined the significance of effects of three transient types, namely that of (i) the presence versus absence of *monocular contrast* transients, (ii) the frequency of *perceptual orientation* transients, and (iii) the presence versus absence of *monocular orientation* transients.

### 2.3 Results & Discussion

We first aimed to test whether the adaptation type in the preceding adaptation phase affected the spatial stability of rivalry in the test phase. Indeed, the fraction mixed percepts during rivalry significantly varied across adaptation types (Figure 2d; repeated measures ANOVA: *F*(4,25) = 11.17, *p* < .001). Qualitative inspection of the pattern of results suggested that the original conflict adaptation condition produced the highest fraction mixed percepts while the conditions with a plaid produced the lowest fraction.

Next, we determined whether we statistically replicated the findings by Said & Heeger (33). While the direction of the effect appeared similar to these previous findings, the conflict and no conflict conditions did not differ significantly according to a two-sided t-test (see Supplementary table 1). A one-sided t-test, which can be argued to be appropriate in case of a prediction based on previous findings, *did* result in a significant effect (*t*(25) = 1.870, *p* = .037). Not surprisingly, the fraction mixed percepts in the first half of the test phase, that is directly after the adaptation phase when effects of adaptation are typically strongest before fading off (39), differed significantly between the conflict and no conflict condition, when tested with a two-sided t-test (*t*(25) = 3.726, *p* = .001).

Next we continued to examine all conditions, including the novel three conditions, in order to determine which transient types drove the adaptation effects. When comparing the patterns of Figure 2c and **2d**, the decrease in perceptual orientations and monocular contrast across conditions matched the decrease in mixed percepts. To explore their individual significance of contribution to the pattern of results, we compared the effects of the presence versus absence of each transient type across conditions on the fraction mixed percepts. The first two conditions were the only conditions that included frequent and repetitive perceptual orientation transients and when pooled together they produced significantly higher fraction mixed percepts than the other three conditions, which included less frequent to no orientation transients (Difference: M = 0.054, SD = 0.051; *t*(25) = 5.389, *p* < .001) The first conflict condition was the only condition that included monocular contrast transients and it produced significantly higher fractions mixed percepts than the other four conditions, which did not include monocular contrast transients (Difference: M = 0.059, SD = 0.075; *t*(25) = 3.986, *p* < .001). The second and third conditions were the only conditions which included monocular orientation transients and they did <u>not</u> produce higher fractions mixed percepts than the other conditions without monocular orientation transients (Difference: M = 0.013, SD = 0.044; *t*(25) = 1.480, *p* = .151). Lastly, the first, second, and fourth conditions were the only conditions which included an orientation conflict between the eyes and they *did* produce higher fractions mixed percepts than the conditions without orientation conflict, but the effect was ~50% weaker than that of perceptual orientation and monocular contrast transients (Difference: M = 0.028, SD = 0.036; *t*(25) = 3.908, *p* = .001).

To summarize the results of Experiment 1, the pattern of destabilization rates across all conditions is best explained by adaptation to both monocular contrast and perceptual orientation transients. Note that the third and fourth conflict rivalry(-plaid) conditions exhibited a conflict between the eyes but produced a lower fraction mixed percept than the first conflict condition. This latter finding cannot be explained by the conflict detector model of Said & Heeger (33) because conflict was clearly present in the rivalry(-plaid) conditions, predicting an increase rather than the observed decrease in the fraction mixed percepts.

Because the manipulations of perceptual transients and monocular contrast (and conflict) were to some degree correlated across conditions, our next goal was to further disentangle the transient types and measure their individual contributions. As such, we continued to test the effects of monocular contrast transients independently from the other transient types in Experiment 2.

## 3. Experiment 2

### 3.1 Introduction

We have learned from Experiment 1 that it is likely that the presence of both perceptual orientation and monocular contrast transients in the adaptation phase disrupted the spatial coherency in binocular rivalry (i.e., increased the fraction mixed percepts) in the subsequent test phase. However, these two transients types co-varied across the conditions of Experiment 1. In Experiment 2 we manipulated the strength of contrast transients in isolation to further assess to what degree they contributed to incoherent perception during rivalry. We took a slightly different approach as compared to Experiment 1 by manipulating the contrast of the rivalling gratings rather than adding distracting information in the other eye. Based on the findings in Experiment 1, we predicted that a low, compared to a high, grating contrast leads to relatively weak monocular contrast transients during adaptation, eventually resulting in relatively weak adaptation and more coherent rivalry, as characterized by less mixed percepts.

### 3.2 Methods

All aspects of the methods were identical to Experiment 1, except for the participants, duration of the rivalry test phase, and adaptation type conditions. A new group of twenty individuals (age: *M =* 21.9, *SD =* 2.3; 14 females) participated in Experiment 2. The rivalry test phase was shortened from 80s to 40s, because of the prominent effects of adaptation in the first 40s. We again included the original two adaptation conditions of Said & Heeger (33) in the conditional design as a reference (see outmost left and right panel in Figure 3a), as well as two novel conditions for which the contrast of tilted gratings were set at 50% and 25% (see second and third panel in Figure 3a). These two conditions specifically affected the degree of perceptual orientation and monocular contrast transients (see dotted and dashed lines in Figure 3b) and, based on the findings in Experiment 1, we predict that the decrease in contrast should weaken adaptation and decrease the fraction mixed percepts (Figure 3c).

**Figure 3.**
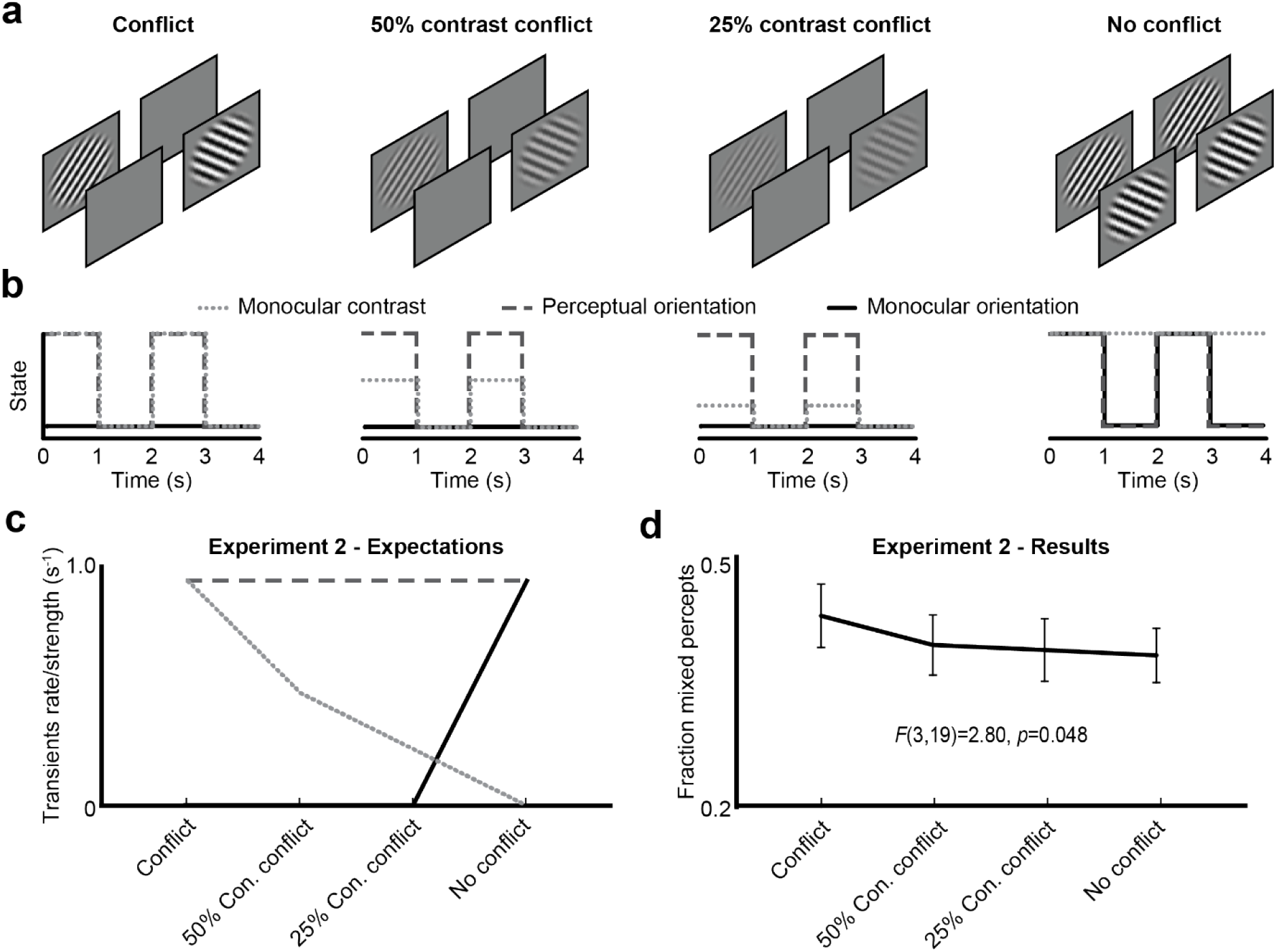
Adaptation conditions, transient profiles, predictions, and results of experiment 2. The design of Experiment 2 contained two original adaptation (first and fourth panel from the left) conditions and two novel conditions (second and third panel) with different time functions of monocular contrast (b). The new conditions varied the strength of monocular contrast transients (c) and the pattern of results followed this manipulation (d).

### 3.1 Results & Discussion

The fraction mixed percepts significantly differed across the four adaptation types (*F*(3,19) = 2.80, *p* = .048), showing a decreasing pattern across conditions (Figure 3d; for post-hoc tests, see Supplementary Table 2). The original conflict adaptation condition produced the highest fraction mixed percepts (M = .43, SD = .17) while the other conditions with a lower grating contrast or no monocular contrast produced significantly lower fractions (M = .39, SD = .15; *t*(19) = 2.529, *p* = .020).

Furthermore, the first conflict condition was the only condition that included 100% monocular contrast transients and it produced significantly higher fractions mixed percepts than the other four conditions (Difference: M = 0.041, SD = 0.073; *t*(19) = 2.529, *p* = .020). The first three conditions were the only conditions which included an orientation conflict between the eyes (and monocular orientation transients) and they did *not* produce higher fractions mixed percepts than the fourth condition without orientation conflict (Difference: M = 0.022, SD = 0.066; *t*(19) = 1.516, *p* = .146).

In sum, we replicated the findings by Said & Heeger and in addition observed that a weaker adaptation contrast decreased the occurrence of mixed percepts during rivalry. The pattern of results of Experiment 2 most closely matched the pattern predicted by the monocular contrast transients, although the flatter and higher pattern than in Experiment 1 suggested that adaptation was again driven by a weighted combination of perceptual orientation transients and monocular contrast transients (i.e., an average of the dotted and dashed line in Figure 3c).

The results of Experiment 1 and 2 together favor a model that combines the effects of perceptual orientation and monocular contrast transients. It remains, however, unclear which of these transient types affects adaptation most. The final Experiment 3 was designed to extract the individual effects of monocular contrast versus perceptual orientation transients.

## 4. Experiment 3

### 4.1 Introduction

Experiment 3 disentangled the effects of monocular contrast transients and perceptual orientation transients by manipulating their presence and absence in opposite manners across conditions.

### 4.2 Methods

All aspects of the methods were identical to Experiment 2, except for the participants and adaptation type conditions. A new group of twenty human individuals (age: *M =* 21.4, *SD =* 2.8; 16 females) participated in experiment 3. The original conflict condition of Said & Heeger again served as a baseline (first panel from the left in Figure 4a) as well as the rivalry-plaid conflict condition from Experiment 1 (second panel in Figure 4a). One novel condition consisted of a rivalry-plaid conflict condition in which the grating’s contrast was lowered by 50% (see third panel in Figure 4a). This manipulation created monocular contrast transients but decreased the frequency of perceived orientation reversals. If both transient types equally strong adapt interocular suppression, both factors should cancel each other and no difference is expected between the full and 50% rivalry-plaid conflict.

**Figure 4.**
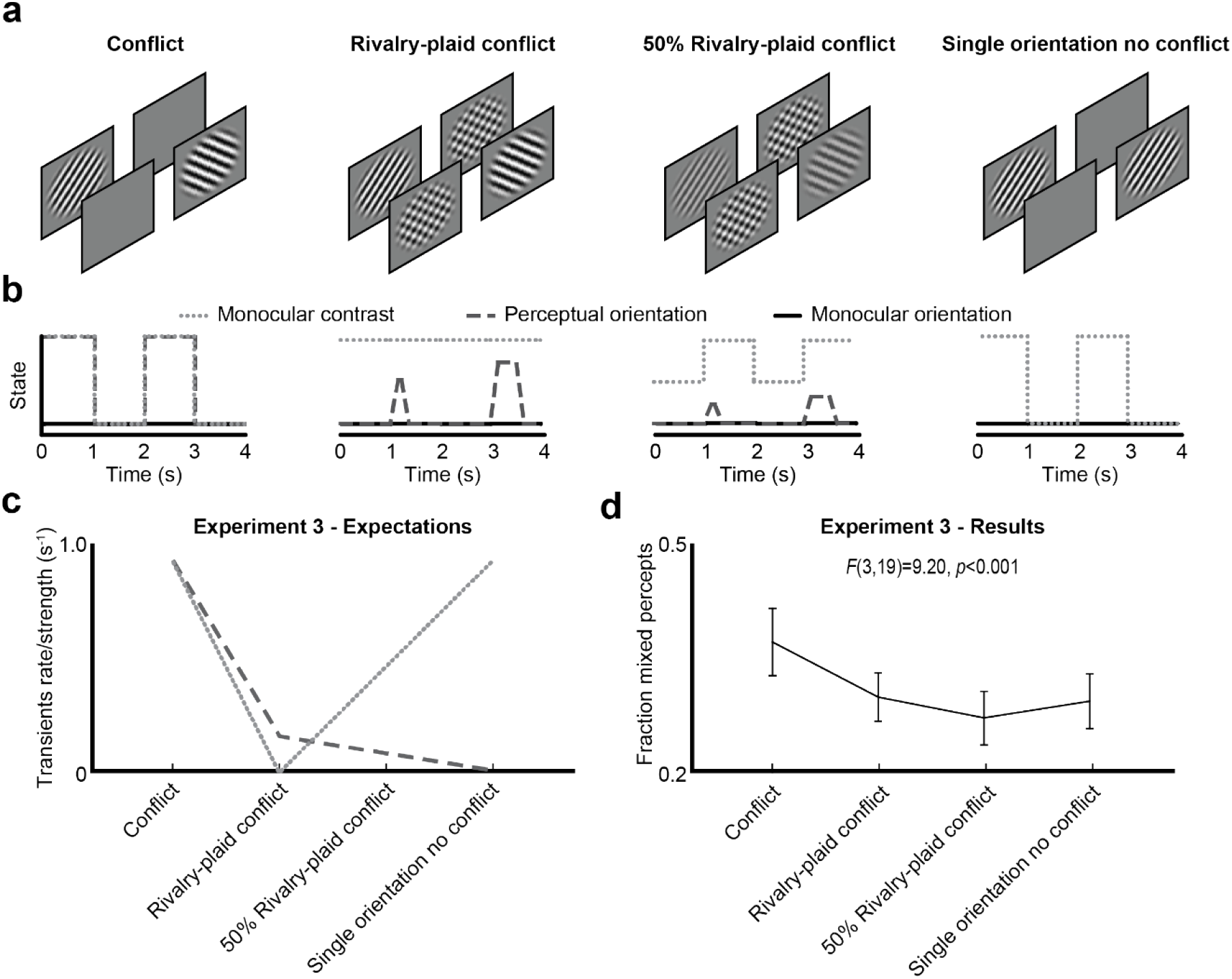
Adaptation conditions, transient profiles, predictions, and results of Experiment 3. Same plots as in **Figure 2** and 3 but now for Experiment 3 with two novel conditions (panel 3-4 at plots (a) and (b)). The pattern of results again reflected a combined weight of monocular contrast and perceptual orientation transients.

We further disentangled the effects of perceptual orientation and monocular contrast transients by solely removing perceptual orientation transients in the last condition (see fourth panel in Figure 4a). This condition consisted of the presentation of a single, non-rotating tilted grating that switched between eyes over time.

The latter three conditions affected the degree of perceptual orientation and monocular contrast transients in opposite manners (see lines in Figure 4b) and each transient type predicted a different pattern of results (Figure 4c).

### 4.1 Results & Discussion

The fraction mixed percepts significantly differed across the four adaptation types (*F*(3,19) = 9.20, *p* < .001), showing a U-shaped pattern across conditions (Figure 4d). The original conflict adaptation condition produced the highest fraction mixed percepts, the rivalry-plaid conflict and single orientation no conflict conditions scored medium fractions, and the 50% rivalry-plaid had the lowest fraction (for post-hoc tests, see Supplementary Table 3). The pattern of results most closely matched a pattern predicted by the combination of perceptual orientation and monocular contrast transients. However, the effects of a weaker perceptual orientation transients and stronger monocular contrast transients in the 50% as compared to 100% rivalry-plaid condition did not cancel each other out. In fact, the 50% contrast rivalry-plaid condition resulted in a significantly lower fraction mixed percepts than the 100% contrast rivalry-plaid condition (*t*(19)=1.787, *p =* .045), indicating that the weakening of perceptual orientation transients had a stronger effect than the strengthening of the monocular contrast transients. In line with this finding, the full removal of perceptual orientation transients with the single orientation no conflict condition decreased the fraction mixed percepts (compared to conflict condition: M = .10, SD = .14) to a similar degree as the removal of half the monocular contrast transients (M = .07, SD = .10; *t*(19) = 1.524, *p* = .144).

Furthermore, the first and fourth condition were the only conditions that included 100% monocular contrast transients and they produced significantly higher fractions mixed percepts than the other two conditions (Difference: M = 0.045, SD = 0.066; *t*(19) = 3.045, *p* = .007). The first condition was the only conditions that included frequent perceptual orientation transients and it produced significantly higher fractions mixed percepts than the other conditions (Difference: M = 0.080, SD = 0.100; *t*(19) = 3.584, *p* = .002). The first three conditions were the only conditions which included an orientation conflict between the eyes and they did *not* produce higher fractions mixed percepts than the condition without orientation conflict (Difference: M = 0.020, SD = 0.048; *t*(19) = 1.817, *p* = .085).

In sum, the results of Experiment 3 suggest that mainly adaptation to perceptual orientation transients and to some extent adaptation to monocular contrast transients cause higher fractions of mixed percepts, indicating more non-exclusive dominance and spatially incoherent rivalry. Note again that almost all conditions included orientation conflict but did not produce similar fractions of mixed percepts. This is in contrast with suggestions by Said & Heeger (33).

## 5. Model – Weighted combinations of transient types

The patterns of results in Experiment 1-3 indicated that the spatial instability of rivalry, measured as the fraction mixed percepts, is most likely enhanced after adaptation to a combination of monocular contrast transients and perceptual orientation transients, but not by monocular orientation transients and not by the presence of orientation conflict between the eyes. To further support this interpretation and to determine the degree of contribution of each individual transient type, we created a step-wise general linear model with the three transient types (monocular contrast, perceptual orientation, and monocular orientation) as well as conflict as predictors of the fraction mixed percepts. The model also included an experiment-dependent intercept α to take into account variance created by differences in the groups of participants across experiments. The fraction mixed percept of the conflict condition, which was included in each experiment, served as the intercept α (Experiment 1: M = .36; Experiment 2: M = .43; Experiment 1: M = .37). The fitted model predicted the results very well, with a root mean squared error (RMSE) of 3% and an *r*^2^ of. 95 (Figure 5). The betas (i.e., slopes) of the factors monocular orientation transients (*β* = 0.004, *p* = .824) and conflict (*β* = 0.005, *p* = .657) were not significant and therefore removed from the model. The final model’s betas for monocular contrast (*β*_c_ = 0.021, *p* = .041), perceptual orientation (*β*_o_ = 0.071, *p* < .001) transients, and experiment-dependent intercept (*β*_g_ = 0.710, *p* = .001) were significant. We conclude from this model that the presence of transients during the adaptation phase, whether produced by a change in grating orientation or contrast, and whether perceived or not, adapted and weakened interocular suppression, and disrupted the spatial coherence of the percept in subsequent binocular rivalry.

**Figure 5.**
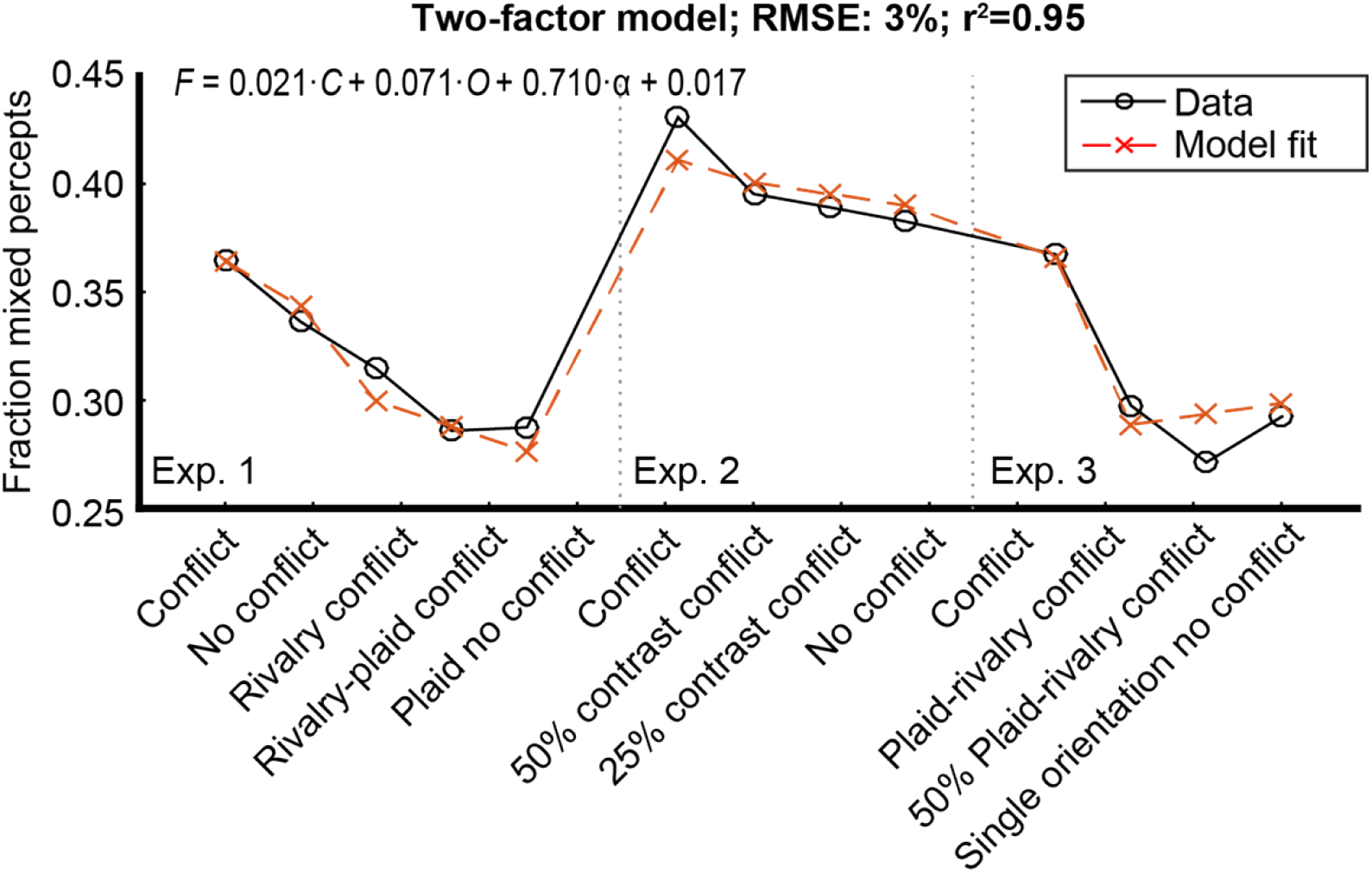
General linear model results. Modelled fraction mixed percepts (dashed red crosses) across the conditions for all experiments as compared to ground truth results (solid black circles) with the factors monocular contrast and perceptual orientation (and an intercept per experiment). The formula is the result of a general linear model with *F* as fraction mixed percepts, *C* as the presence (1) or absence (0) of monocular contrast, *O* as the presence or absence of perceptual orientation transients, and α as the fraction mixed percepts of the conflict condition per experiment (see most left panels in plots (d) in Figure 2–4) that served as an intercept to take into account group differences across experiments.

## 6. General discussion

With a set of three experiments we have assessed whether the precedence of mixed percepts during BR is affected by adaptation to the frequency and strength of stimulus transients or to the degree of interocular conflict as suggested by previous research. The visual transients during adaptation consisted of changes in monocular contrast, perceptual (binocular) orientation, and monocular orientation as a function of time. The fraction mixed percepts, used as a proxy of the degree of the weakening of interocular suppression and spatial destabilization of BR, showed a pattern across a total of 9 distinct conditions that was almost perfectly explained by incidence rates of monocular contrast and perceptual orientation transients. Monocular orientation transients and the presence of a conflict between the eyes as defined in previous work (33) did not explain variance in the pattern of fraction mixed percepts to that degree. We conclude that visual transients affect the depth of interocular suppression during adaptation, resulting in weak, shallow, spatially incoherent binocular rivalry thereafter. Even though monocular contrast transients were inherent to conflict between the eyes in one critical condition (i.e., a blank in one eye and an oriented grating in the other eye), the fact that perceptual orientation transients affected the fraction mixed percepts in the absence of conflict, deems the explanation of visual transients the most parsimonious.

The question remains how transients relate to interocular suppression. We suggest that exogenous, involuntary attention may mediate the link between transients and the adaptation of interocular suppression. Even subtle transients (i.e., cues) to one eye automatically draw attention and can bias perceptual dominance towards that eye (3, 40–44). Similarly, subtle difference between the eyes also attract attention, as demonstrated with a change blindness (45) and visual search paradigm (46–48). As dominance of both eyes is strengthened when attention is drawn to both eyes, the mutual, reciprocal suppression between the eyes is also strengthened (49). Our suggestion therefore is the following: the (visual) transients during the adaptation phase attract attention towards the images and, as a result, increase their mutual inhibition (and thus the amount of interocular suppression). As a result, the strength of mutual inhibition is decreased after adaptation, leading to more shallow rivalry (and hence more mixed percepts) during the following adaptation phase.

An alternative explanation is related to working memory. Sterzer & Rees (50) identified a brain network including parietal and prefrontal areas involved in working memory to become active when dominance in binocular rivalry was temporally stabilized using intermittent blank presentations as strong transients. In line with this knowledge and an initial proposal (35), they suggested that the sudden disappearance of an image during binocular rivalry activates mnemonic processes dedicated to hold the previously seen image in memory and prioritize it for visual awareness the moment it reappears. This memory process is not restrained to only the most recent image but likely holds and biases perception based on images that are observed for at least the last sixty seconds (51). As an image is prioritized, it will also exert stronger suppression to the rivalling image. As the case in the current study, when both images are subject to transients, both will be prioritized and will mutually inhibit each other, that is strengthen interocular suppression and proliferate its adaptation.

It is not unlikely that the effects of working memory and attention on interocular suppression interact. The sudden aspect of transients may (involuntarily and unconsciously) both draw attention and strengthen the (mnemonic) representations of previously seen images, therewith enhancing their inhibitory influence on competing images. However, neither explanation requires adaptation of a specialized conflict detection mechanism. In the model put forth in Said & Heeger, this mechanism is based on the idea of ocular opponency neurons (34, 52, 53). Although such neurons appear likely candidates for involvement in binocular rivalry, and the initial prediction of the model by Said & Heeger that included a conflict detection mechanism explained their data well, the results reported here cannot be unified under that model. As such, we currently see no evidence that mechanisms based on ocular opponency neurons should be included in models of binocular rivalry.

It is important to note that in our study the intermittent presentation of blanks had a stronger effect on adaptation than the intermittent presentation of plaids. A similar effect has been reported before (54), showing that the presentation of interleaved blanks enhanced the temporal stabilization of rivalry more than plaids. As blanks are more distinct from the orthogonal images and therefore more conspicuous, it makes sense that intermittent presentation of blanks adapted interocular suppression stronger than plaids. This conclusion may appear at odds with our observation that the monocular contrast transients (i.e., blanks) disrupted the spatial coherence of rivalry slightly weaker than perceptual orientation transients. Note however that the monocular contrast transients were not visible but the perceptual orientation transients *were* visible to the observer. As the visibility of transients is positively linked to the degree of drawing attention exogenously (55) and the suppressive strength of an evoked traveling dominance wave (26), it is not unexpected that the perceptually visible orientation transients adapted interocular suppression most. Our observation that a relatively high rate of orientation transients (e.g., see rivalry condition) increased the fraction mixed percepts more than a relatively low rate (e.g., see plaid-rivalry condition) further confirms the modulatory effect of transient visibility on the adaptation of interocular suppression. Although out of the scope of the current study, it would be interesting to investigate whether perceptual (and thus visible) contrast transients adapt interocular suppression to a similar degree as the perceptual orientation transients that we investigated here. A useful paradigm to test this would be intermittent presentation in which conflicting gratings disappear and appear as a function of time (35, 56),

To conclude, perceptual stability as expressed in the precedence of mixed percepts and traveling waves during rivalry is weakened when the eyes are stimulated beforehand with many, strong transients. Future work may shed light on the effect of visible and invisible transients on maintaining and adapting to visual representations.

## Additional Information

### Author Contributions

All authors designed the experiments. Author YdK collected the data. MN, SS, and YdK programmed the experiments and analyzed data. Author MN & CP wrote the concept paper and all other authors contributed to the final paper.

## 7. Acknowledgments

Not applicable

## 9. Supplementary materials

**Supplementary Table 1.** Post-hoc, paired, two-tailed t-test comparisons of fraction mixed percepts between conditions for Experiment 1.

| | No conflict<br>($M = 0.34$;<br>$SD = 0.18$) | Rivalry conflict<br>($M = 0.31$;<br>$SD = 0.16$) | Rivalry-plaid<br>($M = 0.29$;<br>$SD = 0.15$) | Plaid no conflict<br>($M = 0.29$;<br>$SD = 0.17$) |
| --- | --- | --- | --- | --- |
| Conflict<br>($M = 0.36$;<br>$SD = 0.18$) | $t = 1.870$,<br>$p = .073$ | $t = 3.291$,<br>$p = .003$ | $t = 4.350$,<br>$p < .001$ | $t = 4.443$,<br>$p < .001$ |
| No conflict | | $t = 1.924$,<br>$p = .066$ | $t = 4.013$,<br>$p < .001$ | $t = 4.088$,<br>$p < .001$ |
| Rivalry conflict | | | $t = 2.231$,<br>$p = .035$ | $t = 1.908$,<br>$p = .068$ |
| Rivalry-plaid | | | | $t = -0.140$,<br>$p = .890$ |

**Supplementary Table 2.** Post-hoc, paired, one-tailed t-test comparisons of fraction mixed percepts between conditions for Experiment 2.

| | 50% Contrast<br>($M = 0.39$;<br>$SD = 0.16$) | 25% Contrast<br>($M = 0.39$;<br>$SD = 0.17$) | No conflict<br>($M = 0.38$;<br>$SD = 0.15$) |
| --- | --- | --- | --- |
| Conflict<br>($M = 0.43$;<br>$SD = 0.17$) | $t = 2.065$,<br>$p = .026$ | $t = 1.874$,<br>$p = .038$ | $t = 2.695$,<br>$p = .007$ |
| 50% Contrast | | $t = 0.433$,<br>$p = .335$ | $t = 0.821$,<br>$p = .211$ |
| 25% Contrast | | | $t = 0.315$,<br>$p = .378$ |

**Supplementary Table 3.** Post-hoc, paired, one-tailed t-test comparisons of fraction mixed percepts between conditions for Experiment 3.

| | 100% Rivalry<br>plaid conflict<br>($M = 0.30$;<br>$SD = 0.14$) | 50% Rivalry<br>plaid conflict<br>($M = 0.27$;<br>$SD = 0.15$) | Single orientation<br>no conflict<br>($M = 0.29$;<br>$SD = 0.15$) |
| --- | --- | --- | --- |
| Conflict<br>($M = 0.37$;<br>$SD = 0.19$) | $t = 2.700$,<br>$p = .007$ | $t = 4.079$,<br>$p < .001$ | $t = 3.446$,<br>$p = .001$ |
| 100% Rivalry-plaid conflict | | $t = 1.787$,<br>$p = .045$ | $t = 0.400$,<br>$p = .347$ |
| 50% Rivalry-plaid conflict | | | $t = -1.524$,<br>$p = .072$ |

